# Forging together: Parallel adaptation to minewater pollution in brown trout (*Salmo trutta* L.)

**DOI:** 10.64898/2026.06.09.731039

**Authors:** D. R. Osmond, J. R. Paris, J. Ferrer Obiol, J. R. Whiting, M.W. Bruford, J. R Stevens

## Abstract

Mining pollution is an important stressor of freshwater communities worldwide. Persistence in these environments requires adaptation, yet identifying the mechanisms responsible in wild systems remains challenging due to differing water chemistries and genetic backgrounds. Parallel evolution presents a powerful framework for identifying adaptive mechanisms through repeated directional change. Here, we seek to understand the mechanisms that enable brown trout (*Salmo trutta*) to survive in metal-polluted rivers. Using low-coverage Whole Genome Resequencing (lc-WGS) we analyse paired metal-polluted and non-polluted brown trout populations across the British Isles. Metal-impacted populations show reduced nucleotide diversity and increased genomic divergence from paired control populations. We observe strong signatures of parallel adaptation in populations with shared geography, emphasising the importance of standing genetic variation in rapid adaptation to pollution. We identify a candidate region of 0.5 Mb on chr25 that shows strong parallel adaptation across multiple pairwise comparisons, including a shared signal among populations experiencing highly divergent water chemistries. The chr25 region contains the genes oestrogen receptor (*esr2b*) and a potassium-gated ion channel (*kcnh5b*), both of which are linked to developmental and osmoregulatory functions known to be disrupted by metals. Using population branch statistics (PBS) and scans for selective sweeps, we also identify population-specific candidate loci, yet putatively selected regions repeatedly converge on shared gene families and functional pathways. Our findings reveal both parallel and unique evolutionary responses to anthropogenic pollution in wild fish, highlighting convergent adaptive pathways to diverse pollutants in teleosts.

## Introduction

As human activities increasingly reshape natural landscapes, many species are confronted with novel and challenging environments characterised by anthropogenic disturbance (Otto, 2018). As a result, organisms remaining *in situ* must rapidly adapt to survive and reproduce in conditions that often differ drastically from their historical environments. Anthropogenic pollution comes in many forms and can comprise a range of more or less significant selective pressures, as species encounter contaminants such as toxic metals (Banks et al., 1997), pharmaceuticals (Milla et al., 2011), sedimentation (Kemp et al., 2011), eutrophication (Weijters et al., 2009) and persistent organic pollutants (El-Shahawi et al., 2010). This is of particular relevance to freshwater ecosystems, which are currently experiencing some of the most rapid declines in biodiversity (Deinet et al., 2020). Aquatic organisms are particularly vulnerable to pollution due to the constant contact of exchange surfaces, e.g. skin, gills, gut, with pollutants dissolved in their environment (Deinet et al., 2020). These environmental stressors can lead to adaptive responses at physiological, behavioural, and genetic levels (Coffin et al., 2022; Hamilton et al., 2017; Jacquin et al., 2020; Oziolor and Matson, 2014; Whitehead et al., 2017). Characterising how species adapt to pollutants in their environment is crucial for understanding the evolutionary impacts of environmental stressors. However, this is often challenging to quantify in natural environments, where exposure typically includes mixtures of compounds with potentially differing additive and non-additive responses. Indeed, in minewater polluted environments, complex mixtures of metals that vary in concentration and bioavailability are likely to impose distinct selective pressures, such that adaptation may reflect responses to the combined effects of multiple stressors rather than to any signal metal in isolation (Das et al., 2016).

At a genetic level, parallel adaptation and convergent evolution provide robust evidence of evolutionary processes driven by environmental pressures (Roesti et al., 2014). Parallel adaptation refers to repeated directional evolutionary change within genomic regions underpinned by the same haplotypes. Convergence relies on a similar evolutionary mechanism at shared loci but differs from parallel adaptation in that different polymorphisms within these shared regions are functionally responsible in each population (Cerca, 2023). Biological replication strengthens evidence of a given stressor in driving adaptation, rather than another biological or environmental factor which may have confounding selective effects (Fraser and Whiting, 2020). Replicate systems are of particular value in understanding the mechanisms and constraints of adaptation to recent anthropogenic stressors, where environmental change is often rapid and there is limited evolutionary time for *de novo* mutations to arise (Lee and Coop, 2017). Recent examples of such evolutionary repeatability include studies of convergent evolutionary responses of sister woodpecker species to local environmental pressures (Moreira and Smith, 2023), adaptation within pest species for insecticide resistance (Cohen et al., 2022), and adaptation to fishing pressure in Atlantic cod populations (Reid et al., 2023). Many insights into the dynamics of repeated adaptation to pollution in fish populations, have come from the study of polychlorinated biphenyls (PCB) adaptation in Atlantic killifish (*Fundulus heteroclitus*), using paired comparisons of pollution-impacted and control clean populations within geographic regions (Reid et al., 2016; Whitehead et al., 2017). As such, replicated systems provide both robust evidence of genomic regions responsible for adaptation and a powerful framework for testing the extent to which this adaptation proceeds through predictable, repeated genomic changes.

Brown trout, *Salmo trutta* (L.), are a widespread teleost fish distributed from Iceland to North Africa, where they occupy a diverse range of habitats (Elliott, 1989). As a result of their ubiquity, brown trout have long been used as a biological model for understanding how different environmental gradients shape genetic diversity and local adaptation (Bekkevold et al., 2020; Ferguson et al., 2019; Kurland et al., 2023; Prodöhl et al., 2017). In particular, brown trout are known to occupy rivers which drain industrially mined regions with high concentrations of dissolved metals, where many other organisms, including other fish species, are extirpated by acute toxicity (Mullinger and Talbot, 2004; O’Grady, 1981). Indeed, this stressor is of high importance to freshwater systems, with recent studies showing that dissolved zinc and copper concentrations are the most significant environmental variables associated with aquatic invertebrate biodiversity (Johnson et al., 2025). Compared to trout from relatively unaffected rivers, metal-impacted trout have highly elevated concentrations of metals in their tissues (Brotheridge et al., 1998; Paris et al., 2024; Uren Webster et al., 2013), show distinct transcriptomic responses to metals both *in situ* (Uren Webster et al., 2013), in gill models (Bury et al., 2014) and in control water (Paris et al., 2024) and are genetically distinct (Osmond et al., 2024; Paris et al., 2015, 2024). However, the importance of genome-wide genetic adaptation to minewater pollution has not been explored.

In this study, we investigate the signatures of parallel and non-parallel genetic adaptation to metal pollution in brown trout sampled from across Britain and Ireland. We use a low-coverage whole genome sequencing approach to examine genome-wide diversity in phylogeographically distinct populations of brown trout exposed to minewater pollution. We apply recently developed methods for detecting genomic regions under parallel adaptation, providing strong evidence for the role of genetic adaptation. We investigate the role of chemical similarity and local shared standing genetic variation in determining parallel and unique evolutionary responses within brown trout populations. Our study assesses whether different mixtures of toxic metals experienced by these fish select for shared mechanisms of adaptation and provides understanding into the constraints of rapid adaptation to anthropogenic stressors.

## Results

Informed by previous observation of repeated patterns of genetic divergence of metal-impacted trout populations across the British Isles (Osmond et al., 2024), we sequenced 154 trout sampled from a total of 19 sites; comprising ten populations from metal-impacted regions, paired with samples from nine control populations from the same regions (Figure 1a). Each metal-impacted site comprises a unique mixture of dissolved metals (Figure 1b), with major axes of chemical similarity separating metal-impacted populations into groups associated with lead/cadmium/zinc toxicity (e.g. CWB - Figure 1c) and copper/aluminium toxicity (e.g. AGA - Figure 1c). We sequenced 137 individuals using lcWGS (∼4x), 12 individuals at 15X and five individuals at 40X (Table 2), resulting in 4,875,231 high-confidence SNPs.

**Figure 1.**
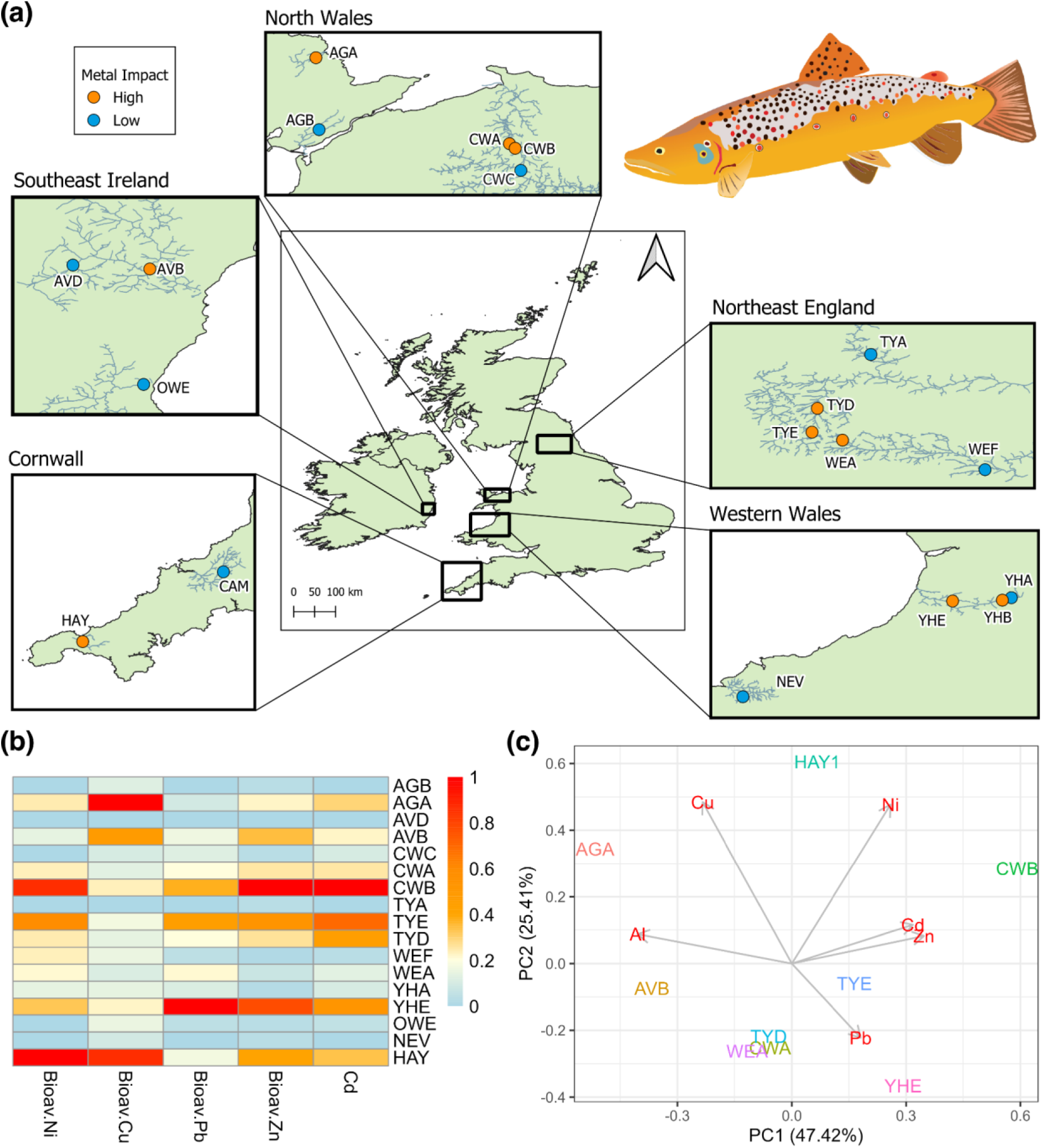
a) Sampling distribution of brown trout (*Salmo trutta*) populations from across the British Isles. Sampling location of metal-impacted populations is given by orange points and relatively unimpacted populations in blue. b) Heatmap of relative metal-impact at sampled sites with bioavailable concentration of nickel, copper, lead, zinc and total dissolved cadmium concentrations have been min-max normalised (0-1). c) Principal Component Analysis (PC1 v PC2) of minewater-impacted populations across axes of relative (min-max normalised) dissolved metal levels.

### Metal impact interacts with phylogeographic history

We first investigated the genetic relationships among sampled populations. Treemix (Pickrell and Pritchard, 2012) analysis revealed high drift for the metal-impacted populations of northeast England (WEA, TYD and TYE), southwest England (HAY) and Anglesey (AGA) (Figure 2a). Credible migration edges were also observed between adjacent populations, including the control populations in northeast England (TYA and WEF) and the metal-impacted populations in north Wales (CWA and CWB) (Figure 2a). Principal Component Analysis showed nested genetic differentiation of metal-impacted and paired control populations within broader geographic patterns of genetic structure (Figure 2b). PC1 (4.81% of the variance) was driven by divergence between populations from northeast England and from the Western British Isles. PC2 (3.40% of the variance) was driven by divergence of the metal-impacted populations Hayle (HAY) and Afon Goch (AGA), with PC3 (2.74% of the variance) driven by divergence of one of the metal-impacted populations from northeast England (TYD) from the other two metal-impacted populations in this geographic region (WEA and TYE) (Figure 2b). The average genome-wide pairwise *F_ST_* between populations ranged from 0.03 (WEF and TYA) to 0.22 (TYD and HAY1) (Figure 2c).

**Figure 2.**
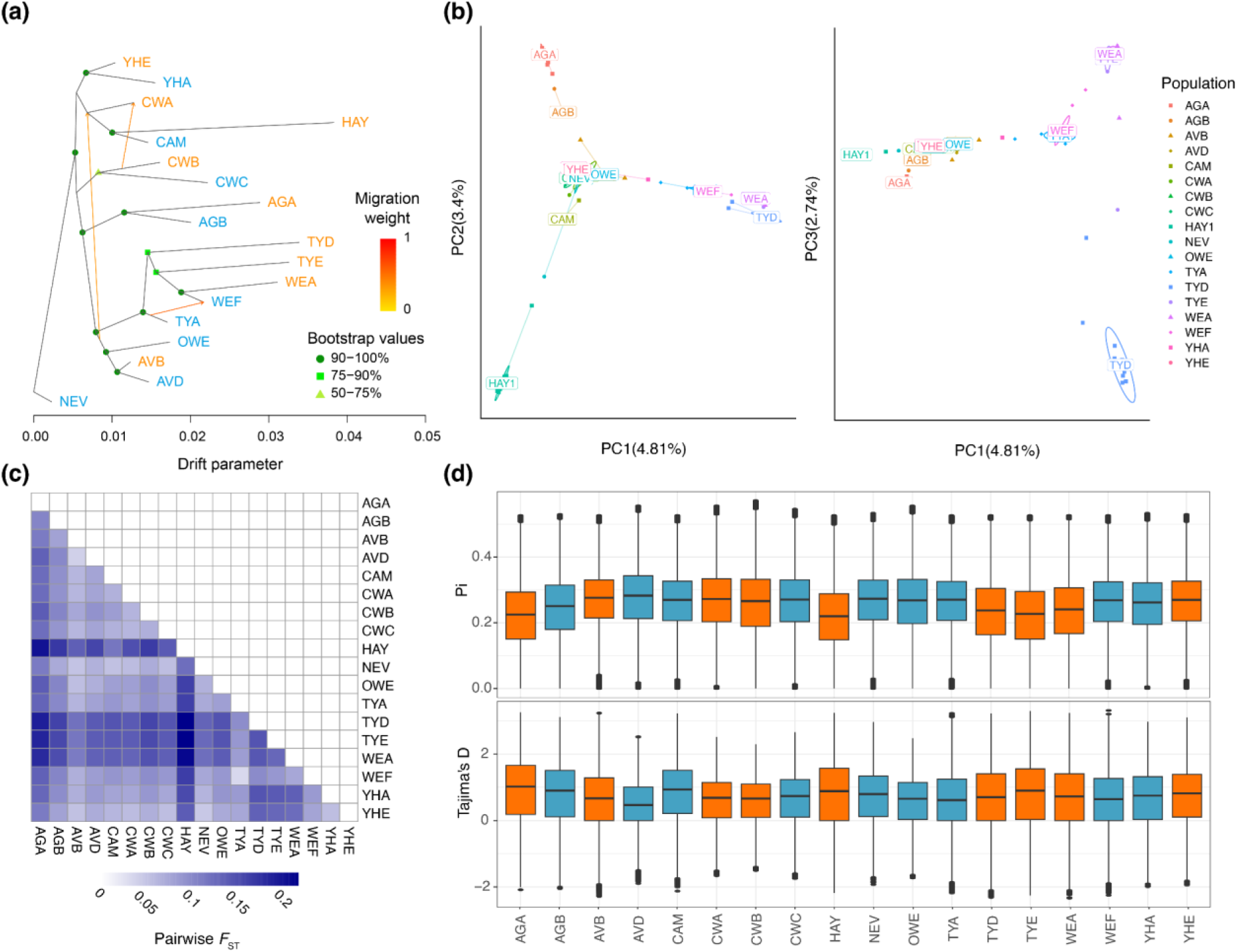
Neutral genetic structure and diversity across the 18 sampled populations of brown Trout from metal-mined regions of Britain and Ireland. a) Treemix population tree of sampled populations, rooted on the river Nevern (NEV) population showing highest likelihood migration edges. Bootstrap support for each node is represented by shape of point and shade as indicated in the key and migration weight indicated by shade of arrow between branches, with higher weight being darker. Minewater impacted populations are labelled in orange, with the relatively clean control populations in blue. b) Principal Component Analysis (PCA) for 149 individual brown trout at 1,993,972 unlinked SNP loci. The x-axis gives the first principal component axis (PC1), representing 4.81% of the variance, with the y-axis in the left plot (PC1 vs PC2) giving the second (PC2) axis (3.40% of variance) and the y-axis in the right plot (PC1 vs PC3) the third (PC3) axis (2.74% of variance). c) Average global pairwise *F_ST_* between each of the sampled populations, with greater genetic distance indicated by darker shading. d) Nucleotide diversity (π) and Tajima’s D in 10 kb windows across the genome for each of the sampled populations. Metal-impacted populations are highlighted in orange, with the relatively clean control populations in blue.

We applied mixed effects models to test the significance of metal-impact on population-level nucleotide diversity (π) and Tajima’s D estimated in 10 kb windows across the genome. Trout populations exposed to the highest concentrations of bioavailable metals consistently showed lower genome-wide π, greater variance in Tajima’s D, and more windows with higher values of Tajima’s D (Figure 2d). There was a significant effect of metal-impact on nucleotide diversity (χ^2^ =5.934, p = 0.0149), with a decrease in nucleotide diversity of 0.0187 associated with high metal impact. We found no significant effect of metal-impact on Tajima’s D (χ^2^ =0.8794, p = 0.348).

### Evidence of parallel selection across a broad geographical range and the influence of geographical proximity and chemical similarity

To investigate regions of parallel genomic change among populations, we utilised AF-vapeR (Whiting et al., 2022). This method quantifies multivariate constraints among genotype change vectors calculated between matched population pairs, in this case clean and metal-impacted pairs. Large eigenvalues on eigenvector 1 imply constraint in haplotypic change across a genomic window that may be indicative of parallel, or antiparallel, allele frequency change among population pairs. Five separate population vectors were applied to examine parallelism across: i) the total dataset from Britain and Ireland; (ii) populations from Wales; iii) populations from northeast England, and among populations exposed to similar mixtures of metals, namely iv) copper-impacted populations, and v) lead-impacted populations.

Across the total dataset comprising all Britain and Ireland population vectors, seven genome windows were identified as significantly parallel (> 99.9%). However, no windows were identified as fully parallel, *i.e.*, parallel windows were also found to be anti-parallel in other population pairs. Using a less-stringent cut-off (> 99%), 74 significant windows were identified, five of which were found to be fully parallel with no anti-parallel signal. Fully-parallel windows were located on chr9 (one window), chr25 (two consecutive windows) and chr28 (two consecutive windows) (Figure 3a).

**Figure 3.**
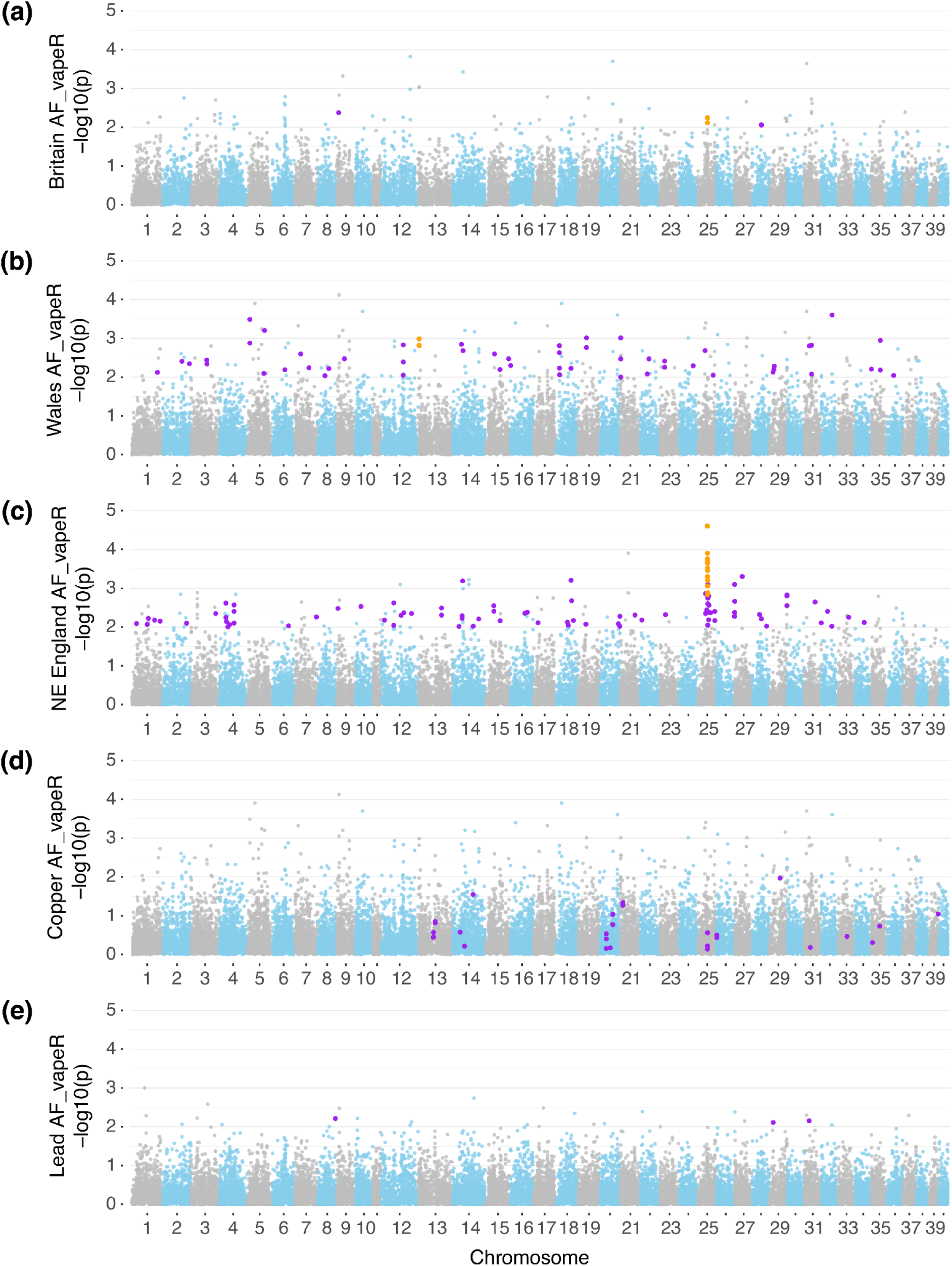
Manhattan plots of AF-vapeR selection scan statistics. The X-axis shows the location of windows across the genome. Chromosomes are alternately shaded grey and blue for visualisation purposes. The Y-axis shows the negative common logarithm of the empirical p-value for each window, with fully parallel significant windows highlighted in purple. Significant parallel windows overlapping candidate regions identified by PicMin are highlighted in orange. Each panel represents a different set of population vectors, representing the full geographic distribution across: (a) Britain and Ireland; (b), populations from Wales; (c) northeast (NE) England; (d) copper-impacted populations; and (e) and lead-impacted populations.

Within Wales, 19 significant (>99.9%) windows were identified along the first eigenvector; of these, three windows had no anti-parallel population pairs and were found on chr1 and chr27 (Figure 3b). A total of 184 windows had eigenvalues > 99% of null permutations along the first eigenvector, of which 56 had no anti-parallel populations. Two of these fully parallel windows were situated within a region in chr25 that showed a large cluster of windows (21 fully parallel windows from 24.35-26.17 Mb) in the northeast England analysis (Figure 3c). In the latter, AF-vapeR identified 29 significant windows (>99.9%), 25 of which were fully parallel in all three population pairs. The large region on chr25 was identified as being under significant parallelism in all population pairs. Other parallel windows were identified on chr26 and chr27 (Figure 3c).

Within the vector set of predominantly copper-impacted rivers and their clean comparators, nine significant windows were found along the first eigenvector using a cut-off >99.9%, two of which were fully parallel and adjacent to each other at approximately 35 Mb on chr20. At a significance cut-off of >99%, we recovered 78 windows, 27 of which were fully parallel in all population pairs (Figure 3d). For the analysis of the lead-impacted populations, only one significant window (not fully parallel) was identified using a cut-off >99.9% along the first eigenvector. At a cut-off of >99%, there were 28 significant windows, of which three were fully parallel (Figure 3e).

Taken together, these results give limited evidence for large parallel haplotype re-use in driving shared adaptive responses to metal-pollution. Nevertheless, a candidate region on chr25 emerged consistently across analyses: first with two consecutive fully parallel windows within the broad total-dataset analysis and again identified by multiple significant, fully parallel windows in analyses of population vectors from northeast England and Wales. Notably, geographically localised analyses revealed a greater number of significant parallel windows than the broad analysis, suggesting that parallel haplotype re-use may be more relevant at finer geographic scales. By contrast, greater chemical similarity among population pairs did not correspond to greater detection of parallel haplotype change in the lead and copper-driven populations analyses (Figure 3).

### Evidence for convergent evolution within genomic regions to metal-pollution

To explore convergent evolution without assuming parallel haplotype re-use across populations, we applied PicMin (Booker et al., 2023). This method uses the theory of order statistics to evaluate the likelihood of the nth smallest p-value for each genomic window across population pairs, enabling the detection of significant repeatability without predefined significance thresholds or an assumption of shared haplotypes within divergent regions. Across a balanced vector of populations from the broad geographical dataset, we identified a total of 170 significant (FDR = 0.5) gene-overlapping windows encompassing 306 individual genes in two or more population pairs. Using a more stringent cut-off (FDR = 0.2), we retained seven significant overlapping windows, which together contained 11 protein-coding genes. These seven windows included a region on chr1 at 60.1 Mb, which overlapped with the potassium-gated ion channel (*kcnh6a*) gene, which was driven by convergence across all population pairs. In addition, we identified three windows on chr3, which included the genes *pcdh1b* and *aff2.* Finally, two of the three windows detected on chr25 overlapped with the 24.8 Mb region also identified by AF-vapeR. The two most significant windows on chr25 were supported by signals of convergence across four and five population-pair comparisons, respectively. Furthermore, all population pairs except the Anglesey pair showed significantly elevated *F_ST_* within this region. The third outlier window on chr25 was driven by strong convergence at two population pairs (Cornwall and Tyne), which included the gene fibroblast growth factor receptor 3 (*fgfr3*) (Figure 4).

**Figure 4.**
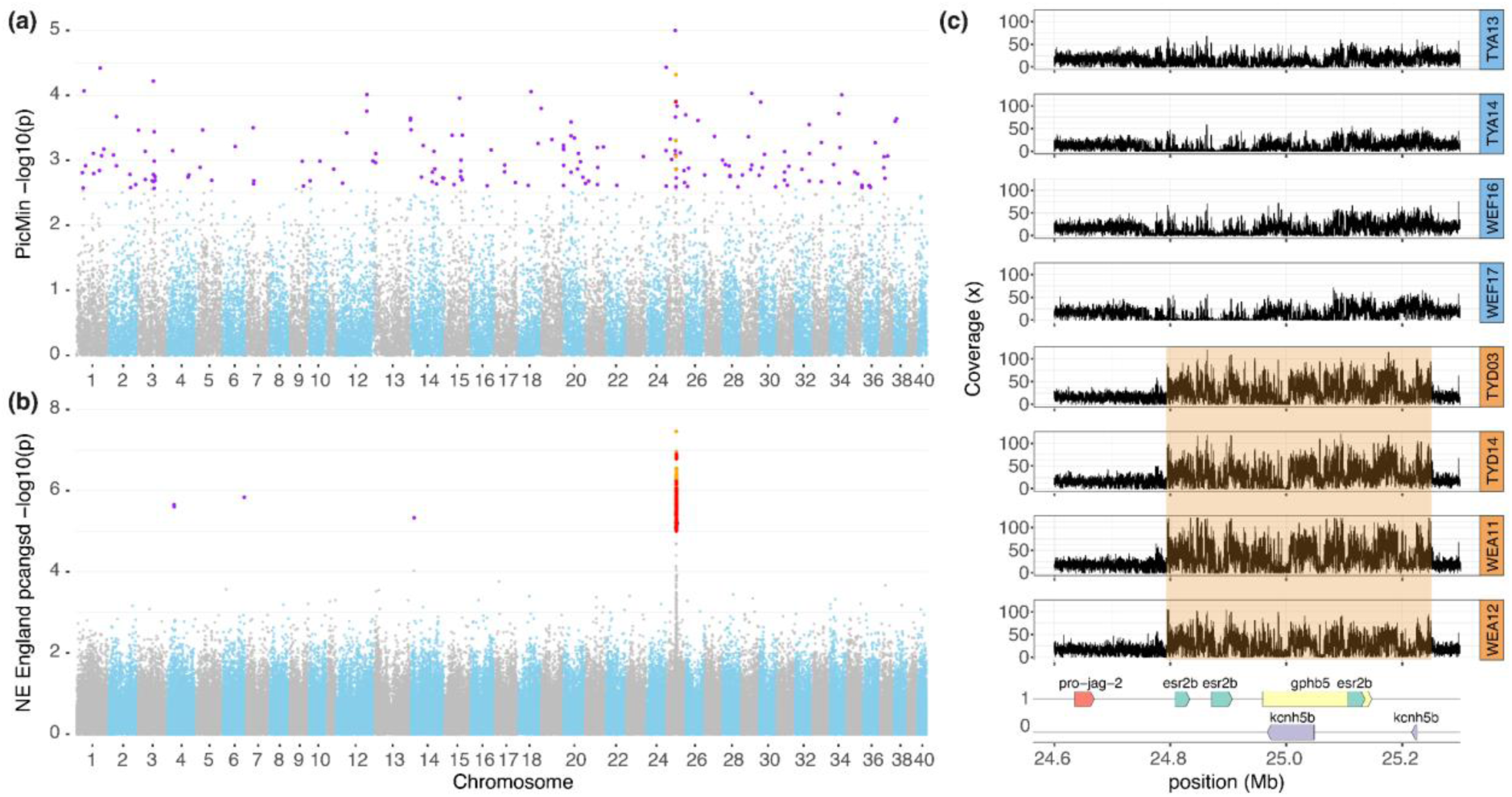
Manhattan plots of selection scan statistics for (a) PicMin analyses of all regional paired trout populations and (b) PCAngsd selection. Candidate regions identified by each method are plotted in purple, with regions supported by two parallel outlier scan methods highlighted in orange and regions supported by all three parallel outlier scan methods (AF-vapeR, PicMin and PCAngsd - overlapping candidates for these limited to northeast England population analyses) highlighted in red. c) Read depth mapping for the 15x sequenced individuals from northeast England around the parallel candidate region identified by all three selection methods (chr25: 24.8-25.1 Mb). Read depth gives support for the potential role of copy number variation in the observed genomic differentiation within this region. The candidate region is highlighted in orange within the metal-adapted individuals. Genes within the region are labelled beneath the plot.

Both methods used for detecting repeated evolutionary change (AF-vapeR and PicMin) detected highly significant windows on chr25, providing independent support for this region as a candidate target of adaptation to metal adaptation. Using PicMin, we further identified additional candidate regions under broad repeated selection, such as a window containing the potassium-gated ion channel (*kcnh6a*) gene, which were undetected by AF-vapeR. This suggests that convergent adaptation to metal pollution may be mediated by selection acting on different haplotypes within shared genomic regions.

### PCA-based selection scans corroborate chr25 region as a candidate for parallel adaptation to metal pollution

Previous studies have suggested that PCA-based methods integrating a probabilistic framework are a powerful method for identifying candidate loci for selection (Lou et al., 2020). To further explore candidate regions and compare the power of these lc-WGS methods with results obtained from AF-vapeR and PicMin, we applied the selection utility within PCAngsd (Meisner et al., 2021) to three groups: i) all individuals in the dataset, and separate analyses using only those sampling sites representing ii) northeast England, and iii) Wales.

The PCAngsd selection analysis with the full Britain and Ireland dataset and with the Wales dataset were not informative for detecting parallel evolutionary regions associated with environmental stressors, due to the principal axes not separating populations along a vector of metal-pollution. However, using the northeast England dataset, the third PC axis (5.94% of the variance) characterised a split between the metal-impacted (TYD, TYE and WEA) and the control (TYA and WEF) populations. Along PC3, one significant outlier window was identified with a highly significant peak in the chr25 region previously detected as an outlier in the AF-vapeR analysis (around 24.8 Mb) (Figure 4a). This region is in close proximity to three homologs of oestrogen receptor 2b (*esr2b*) and two potassium-gated ion channel (*kcnh5b*) genes.

### Evidence of unique evolutionary responses to metal-pollution within individual populations

To identify potential regions indicative of unique adaptation to metal pollutants in each of the populations, we applied two methods with differing assumptions. First, we used population branch statistics (PBS), which compare the target population with a local paired comparison and a third outgroup, to identify windows where the *F_ST_* of the target population is elevated above the genome-wide average (Yi et al., 2010). In addition, we used Sweepfinder2 (DeGiorgio et al., 2016) to identify regions of the genome associated with selective sweeps. We intersected candidates identified from both of these methods to identify genomic regions associated with high *F_ST_*between metal-impacted and control populations, where the site frequency spectrum (SFS) differs from the null expectation.

Across all PBS analyses, we identified 1,272 outlier windows with an empPval <0.005 in at least one population. The majority of these windows were only significant in a single PBS analysis. However, the chr25 region at approximately 24.4-25.2 Mb was identified as an outlier in all three northeast England PBS analyses. Across the metal-impacted populations that were analysed, SweepFinder2 identified 652 significant regions in HAY, 586 in YHE, 447 in AVB, 405 in WEA, 404 in CWB, 323 in TYD, 269 in AGA, and 209 in TYE. After the exclusion of those regions where overlapping outliers were identified in paired control populations, 651 regions were retained in HAY, 482 in YHE, 387 in AVB, 338 in WEA, 331 in CWB, 273 in TYD, 243 in AGA, and 170 in TYE. Overlapping significant SweepFinder2 windows with candidate windows identified in the PBS analysis, we identified 25 overlapping regions in HAY, 18 in AGA, 17 in AVB, 17 in TYD, 16 in CWB, 14 in YHE, 13 in TYE, and 10 in WEA. Gene Ontology (GO) enrichment analyses in these outlier windows (Table 1) showed significantly overrepresented biological processes involved in calcium ion transport, ion transmembrane transport and calcium ion homeostasis, and overrepresented molecular functions encompassing copper ion binding and ion channel activity. A complete list of outlier regions, gene annotations and Manhattan plots can be found in Supplementary Materials 1 and 2.

**Table 1.**
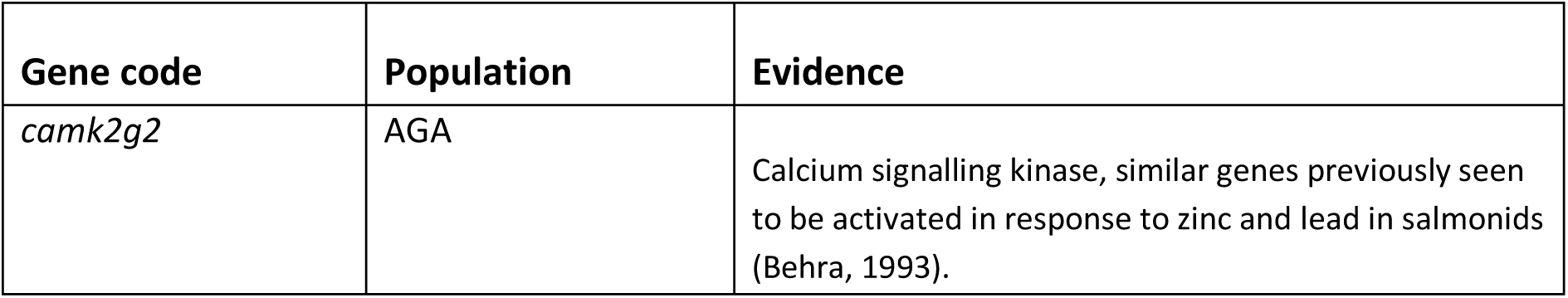
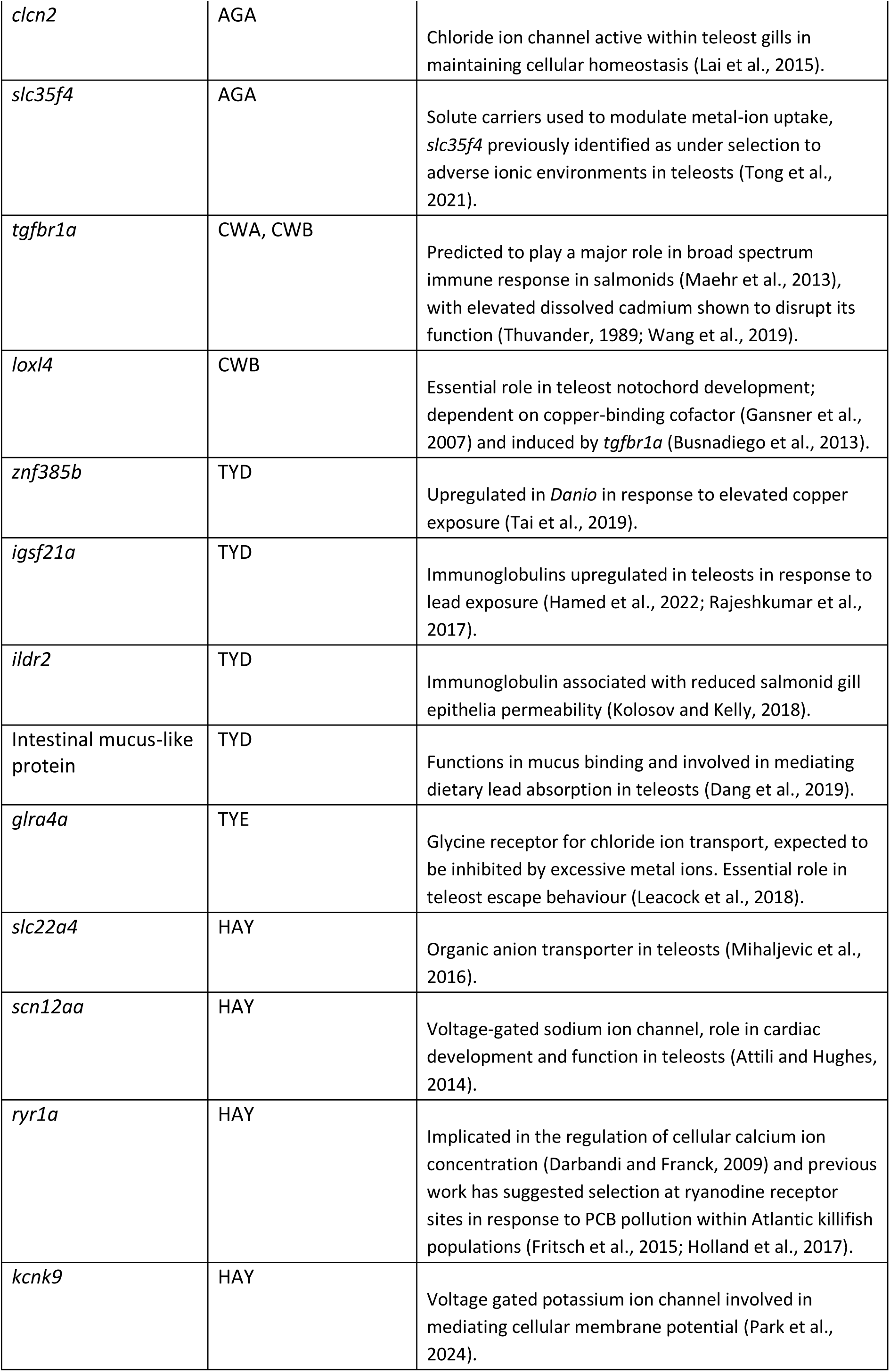
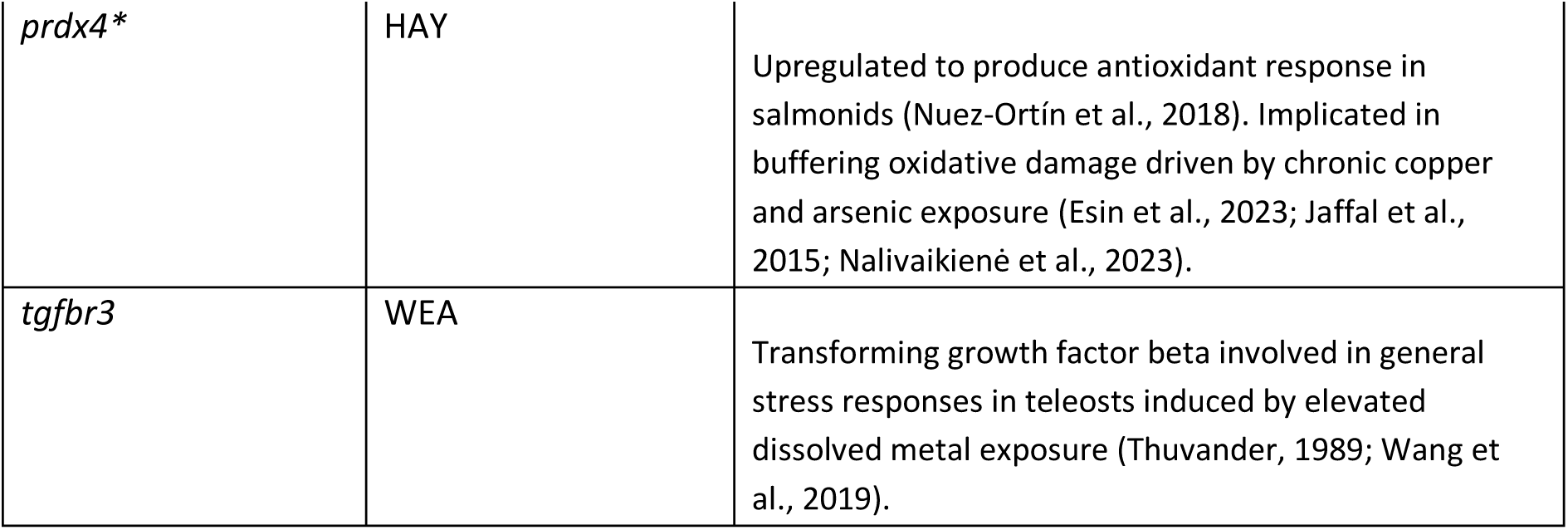
Candidate genes for adaptation to metal-polluted environments observed within individual populations. Unless specified, all loci were found to be under selection by both Sweepfinder2 and PBS analyses, genes marked with * were found to be under selection by PBS and a previous transcriptomic analysis (Paris et al., 2024).

### Connecting evidence of adaptation within genomic regions across multiple analyses

Across AF-vapeR, PicMin and the PBS analyses covering the Britain and Ireland vector set, we uncovered three genes in regions identified as being under full parallelism: *gphb5* (ENSSTUG00000040246), *kcnh5b* (ENSSTUG00000040255) and *esr2b* (ENSSTUG00000040315) (Figure 4c). These genes are all located within a region on chr25 spanning approximately 24.9–25.1 Mb and were among the most strongly differentiated loci in the population vectors representing northeast England. Within northeast England, seven genes were repeatedly identified by all three outlier methods, spanning a broader region on chr25 (24.4–25.6 Mb) that encompassed the candidate region detected in the Britain-wide vector set. Within Wales, two genes were identified in the overlapping windows identified by PicMin and AF-vapeR: *nexmif* (ENSSTUG00000046608) at 39.0 Mb on chr15 and *col15a1b* (ENSSTUG00000035162) on chr31 at 19.4 Mb. This latter region was also associated with a particularly strong putative selective sweep identified by SweepFinder and a PBS outlier in the CWB population from Conwy (northwest Wales), which contains two ion transport proteins (*kcnh2a* and *slc127a*) within 200 kb upstream of the candidate window. In the Wales analysis, we identified 27 overlapping windows between the PBS and AF-vapeR analysis, containing a total of 61 genes. However, no significant outlier windows were found to overlap across all three methods (AF-vapeR, PicMin and PCAngsd). Investigating parallelism by water chemistry, seven genes were identified in the overlapping significant selection windows shared by both PicMin and AF-vapeR in the copper analysis. These included *gphb5*, *kcnh5b, esr2b* and *ppp2r5eb* (ENSSTUG00000042918) spanning the 24.9 –25.3 Mb region of chr25*, atg16I1* (ENSSTUG00000019822) and two protein-coding genes of unknown function around the 0.9 Mb region of chr26. No concordance was observed between any of the selection methods within the vector set including lead-impacted populations.

Structural variants can facilitate adaptation by reducing local recombination and maintaining favourable combinations of alleles, making them important drivers of repeated evolutionary change (An et al., 2024; Li et al., 2024). Calculation and visualisation of read depth at the candidate chr25 region revealed increased read depth within the metal impacted populations of Northeast England (TYD and WEA) compared to relatively clean paired control individuals sequenced at medium - high depth (TYA and WEF) (Figure 4c). Homologous regions of the trout genome exhibited typical consistent read-depth mapping, suggesting that copy number variation within this region may be responsible for the observed genomic divergence between populations.

### Candidate genes for metal-adaptation within the Hayle are supported by transcriptomic evidence

A previous study identified candidate genes which were significantly constitutively expressed in gill and liver tissues in metal-impacted trout after 11 days in metal-free water (hereafter referred to as gill-expressed and liver-expressed genes) (see Paris et al., 2024 for full details). We intersected these expression candidates with the outlier genes identified in this study. Intersection between gill-expressed genes and PicMin FDR < 0.5 genes identified 13 shared genes, including the potassium channel protein *kcnk2b*. No overlap was found with the liver-expressed genes. One overlapping gene was identified in the gill-expressed genes and the Hayle SweepFinder-PBS intersect outlier windows: ryanodine receptor 1a (*ryr1a*), but no overlap was found between these genes and the liver-expressed genes. Ten overlapping genes were found between the PBS Hayle p < 0.005 windows and the gill-expressed genes, and two genes between the Hayle PBS outliers genes and liver-expressed genes, including the peroxiredoxin 4 (*prdx4*) gene. No overlapping genes were identified across gill and liver expression outliers and significant, fully parallel AF-vapeR outliers, in both the Britain and Ireland-wide, and copper population vectors.

## Discussion

Our results demonstrate the role of legacy pollution in shaping genomic diversity and in driving parallel and non-parallel adaptation in metal-impacted brown trout populations. We observed reduced nucleotide diversity in impacted populations, indicative of recent population contractions. We also identified signatures of increased genetic drift in populations most impacted by metal pollution and strong genetic structure between these populations and their control population counterparts. We observed a strong signature of parallel selection in several genomic regions; this was particularly evident in trout populations within the same geographical region. We identified one genomic region as consistently under parallel selection on chr25, even in highly geographically divergent populations across the British Isles. This region was shown to contain genes relevant to mediating metal toxicity, suggesting parallel adaptation to minewater pollution in impacted populations of *Salmo trutta*.

### Metal-impacted trout populations are consistently genetically distinct and show reduced genetic diversity

Across our population structure results, we observed strong genetic structure between paired metal-impacted and relatively unaffected control populations. Analysis of paired metal-impacted and control populations generally resulted in high *F_ST_* values and we observed long branch lengths in our treemix analyses, suggesting metal-impacted populations have experienced high genetic drift. This pattern of nested genetic differentiation between metal and clean populations within geographic regions accords with the patterns observed using a panel of 95 neutral SNP loci (Osmond et al., 2024). We also observed that metal-impacted populations also had a more positively skewed distribution of Tajima’s D and a more negative distribution of nucleotide diversity. Higher average Tajima’s D is typical of populations which have recently experienced a bottleneck, where low frequency polymorphisms have been purged (Gattepaille et al., 2013) and reduced nucleotide diversity suggests that alongside bottlenecks, reproductive isolation or positive selection may also credibly be responsible for shaping the divergence of these populations (Therkildsen et al., 2019). Together, these results highlight the impact of pollution on neutral genetic diversity, with exposed populations, despite persisting in the present environment, being vulnerable through the erosion of genetic variation to future environmental changes (Santos et al., 2013; Švara et al., 2022).

### Evidence of parallel adaptation to a common stressor of minewater pollution

Within our AF-vapeR analysis of the British Isles-wide sample set, we detected relatively few significant outlier windows under parallel selection in all population pairs, and of these, only two adjacent windows on chr25 contained annotated protein-coding genes. When analyses were restricted to vector sets representing smaller geographic regions (*i.e.*, NE England, Wales) a larger number of significant windows were identified as fully parallel. This result could be a function of similarity in water chemistry in the NE England analysis, with lead and zinc being the more common acutely toxic metals within these rivers. Conversely, the analysis for Wales includes populations from highly dissimilar metal pollutants, ranging from the copper-driven chemical environment of the Afon Goch, to the lead, zinc and cadmium-dominated pollution in the Ystwyth and Conwy.

Previous studies have implicated the role of standing genetic variation in rapid adaptation of fishes, such as the selection for the Omy5 region in isolated rainbow trout (*Oncorhynchus mykiss*) populations (Pearse et al., 2014) and hard selective sweeps for parallel adaptation to acidic environments in stickleback (*Gasterosteus aculeatus*; (Haenel et al., 2019). The putatively parallel adaptive genes identified in Welsh trout populations are strongly associated with physiological tolerance to disrupted homeostasis. These include multiple ion-transport and channel genes under parallel selection, including the voltage-dependent calcium channel genes *cacna1aa* and *cacna1ia*, the potassium channel genes *kcnk1b*, *kcnh1a*, and *kcnh2a*, and the anion-channel gene *vdac1*. Consistent with this, GO enrichment analyses revealed significant overrepresentation of ion-channel and ion-binding functions. The disruption of ionic homeostasis of Na^+^ and Ca^2+^ has been demonstrated to be a key mechanism of acute toxicity in salmonids exposed to metals, including cadmium, zinc and copper (McGeer et al., 2000) with inhibition and competition at gill ion transporters (Birceanu et al., 2008; Niyogi et al., 2015). As expected, population-specific signatures were more numerous than shared candidates. However, our results provide evidence for parallel evolutionary responses mediated by shared standing genetic variation among populations exposed to distinct metal mixtures associated with minewater contamination. These convergent responses likely reflect adaptation to the common homeostatic stress imposed by metal pollution.

### Major pathways of metal adaptation and a candidate adaptive region on chr25

A strong candidate locus for parallel adaptation was detected by all methods on chr25, at approximately 24.7 - 25.2 Mb, across a sample set representing populations from all regions examined. The strongest signal of differentiation at this locus was observed within the comparisons of metal-impacted populations from NE England (TYD, TYE and WEA) where the region was independently identified by PicMin, AF-vapeR and pcangsd analyses. Patterns of mapping depth across this region further suggest that the inferred allele frequency divergence may be attributable to underlying copy number variation, implicating structural variation as a potential mechanism of adaptation. Structural variation is increasingly recognised as playing an important role in adaptation (An et al., 2024; Ho et al., 2020; Li et al., 2024; Stenløkk, 2023), including in parallel responses (Battlay et al., 2025; Zong et al., 2021). However, the limited ability of short-read sequencing to span large structural variants constrains our inference of the precise adaptive architecture within this region. Future studies harnessing long-read sequencing will be key for resolving the functional and structural basis of adaptation at this locus.

The chr25 candidate region contains a voltage-gated potassium ion transporter gene (*kcnh5b*) and duplicated copies of the *esr2b* oestrogen receptor gene. Previous research has identified *esr2b* as a target of parallel selection in Atlantic killifish populations exposed to PCB pollution (Whitehead et al., 2017). In addition, several potassium-gated ion channels have also been implicated as under selection within PCB-impacted Atlantic killifish populations (Osterberg et al., 2018). *esr2b* is responsible for gonad development and reproduction (Yan et al., 2019), and its function is known to be disrupted by both cadmium and lead exposure (Chakraborty, 2021; Łuszczek-Trojnar et al., 2014; Vetillard and Bailhache, 2005). Putative selection at this gene may therefore constitute a shared mechanism for buffering the disruptive effects of pollutants on reproductive function, with parallel signatures of selection observed in response to both metal pollution in this study and PCB contamination in Atlantic killifish populations (Whitehead et al., 2017). In addition, exposure to elevated dissolved lead in fish has been shown to have highly disruptive effects on ionic homeostasis across the gills, with lead ions binding to ionic transporters and stimulating upregulation of expression of ion transport pathways (Curcio et al., 2022; Ribeiro et al., 2014). Selection on *kcnh5b* is therefore a likely result of this ionic challenge from lead pollution, with several other ion transport genes also found to be under selection across our parallel selection analyses.

### Limited evidence for chemically similar environments in driving more parallel evolutionary responses

Our environmental PCA and heatmap showed two main groups of metal mixtures to be associated with our sampling sites: (i) Cu and Al, and (ii) Pb, Cd, Zn and Ni. We therefore selected populations representing these two major groups to assess parallelism among populations exposed to broadly similar chemical environments, which we termed the copper and lead analysis vectors, respectively. AF-vapeR analysis of these vectors revealed relatively few candidate genes compared with geographical analyses. Within our analysis of the copper group, only the region on chr25, already identified in the full regional analysis, overlapped with the significant PicMin outliers. In the lead analysis, we found no overlap of genomic windows between the selection outlier tests, with relatively few fully parallel candidate windows above the 99% cut-off in AF-vapeR and no fully parallel outliers >99.9%. The lack of significant outlier windows along these environmental vectors could be due to several reasons: 1) there is a lack of parallel adaptation specific to particular metal mixtures across broad geographical regions; 2) we have an insufficient number of population-pairs within our vector set to provide statistical power to distinguish the signal of parallel adaptation from noise; 3) our normalisation of metal bioavailability and average environmental measures of dissolved concentration does not reliably measure biological relevance. Given the observed parallel adaptation within vector sets of 3-4 comparison pairs in the geographical region analyses, the second of these explanations seems unlikely. Plotting the shared windows among populations in the PicMin FDR < 0.5 outliers (Supplementary Materials 1, Figure S2) does not appear to reveal any clear patterns of shared genomic windows as a function of geographical proximity nor along axes of individual or mixtures of metals. Repeated reductions in genomic diversity associated with metal pollution across populations are also expected to limit the shared standing genetic variation available for selection to generate parallel evolutionary responses among disparate populations. Taken together with the observed parallelism within geographic regions, this suggests a greater influence of shared standing variation within populations than similarity of specific acutely toxic metal mixtures in driving parallel evolutionary responses.

### Evidence of unique adaptation to metal pollution within individual populations

Combined SweepFinder and PBS overlap analyses revealed extensive population-specific signatures of selection, yet these repeatedly implicated shared gene families and functional pathways. Broad-spectrum immune function genes, including members of the TGF-Ꞵ family, were repeatedly identified across populations, consistent with previous evidence that elevated metal concentrations activate these pathways (Thuvander, 1989; Wang et al., 2019). Similarly, these analyses revealed many transmembrane channels involved in maintaining ionic balance across metal-impacted populations. It is known that in environments with elevated dissolved metals, these ions can interact competitively and have deleterious impact on homeostasis (Kwong, 2024). Further functional validation is required to elucidate the role of these candidate genes in individual-population adaptation to metals. However, overlap with transcriptomic evidence from a previous study (Paris et al., 2024) strengthens support for their adaptive relevance. Overall, these results support recent research that suggests selection acts on different genes with shared biological function resulting in convergent trait adaptation despite an apparent absence of genomic parallelism (Martin et al., 2024; Whiting et al., 2024). Notably, ryanodine receptor sites have been identified previously to be under selection in response to PCB pollution within Atlantic killifish populations (Fritsch et al., 2015; Holland et al., 2017), suggesting broad teleost-wide adaptation pathways to diverse chemical pollutants.

Despite chronic exposure to similar mixtures of cadmium, lead, and zinc, and their shared location within the Tyne catchment (northeast England), the West Allen (TYD) and Nenthead (TYE) populations are highly genetically distinct, with this divergence extending to population-specific signatures of adaptation. Within TYE, we found only one instance of positive selection overlapping with PBS outliers with a credible role in adaptation to metal pollution (*glra4a*), which contrasts with several candidate loci identified in the less acutely polluted TYD. The severity of acute metal toxicity at TYE suggests that this population should be facing strong selective pressure and, as such, we might expect to detect more outlier loci. However, the nucleotide diversity of this population was the lowest detected from amongst the NE England sites and was the lowest across our study. The natural isolation of the TYE population above impassable waterfalls may explain both the reduced nucleotide diversity of this population and the apparent reduced adaptive response compared with TYD, as isolated, small populations are known to have reduced adaptive potential (Frankham, 2015; Pavlova et al., 2017). Although stringent criteria were applied to identify candidates supported by overlapping SweepFinder and PBS outliers, transcriptomic support for *prdx4* in the Hayle population, despite the absence of a significant deviation in the site frequency spectrum, suggests that candidate loci may be overlooked when adaptation is assumed to proceed primarily through hard selective sweeps. This is supported by recent studies that suggest that soft, polygenetic adaptation is more frequently associated with rapid adaptation to environmental change (Garud et al., 2021).

## Conclusions

Our results provide compelling evidence for repeated adaptation to minewater pollution in phylogeographically disparate populations of brown trout across the British Isles. Metal-impacted populations were characterised by reduced genetic diversity and increased genetic differentiation from nearby control populations sampled from unpolluted rivers. We identified a diverse set of candidate loci associated with adaptation to metal pollution that were unique to each impacted population, with several candidates additionally supported by transcriptomic evidence, suggesting widespread population-specific genomic responses to chronic environmental contamination. We identified loci repeatedly under selection in multiple comparisons of metal-impacted and control populations, with a candidate region on chr25 showing strong signatures of parallel adaptation in all our population comparisons across multiple analyses. Stronger signatures of parallel adaptation were observed among more closely related populations, highlighting the importance of shared standing genetic variation and potential ongoing gene flow in facilitating adaptive responses and maintaining adaptive potential. Our results therefore suggest that restoring connectivity among isolated populations could play a key role in sustaining genetic diversity and preserving adaptive potential under increasing environmental stress.

## Methods

### Study sites and sampling

Based on patterns of genetic structure and diversity identified in the Osmond et al., (2024) SNP study, we selected a geographically representative subset of populations to provide regional controls and investigate adaptation to metal pollution. To examine water chemistry similarity of sampled sites along multiple axes of different dissolved metals, data were obtained from Natural Resources Wales, the Environment Agency (UK), and Environment Protection Agency (Ireland). The bioavailability of four metals: copper (Cu), lead (Pb), zinc (Zn) and nickel (Ni) were calculated using the Bio-met bioavailability tool v5.0 (bio-met.net/) and these, along with the dissolved concentrations of aluminium (Al) and cadmium (Cd), were individually min-max normalised across the metal-impacted sites. These metals were chosen on the basis of toxicity, availability of data across the sampling sites, and bioavailability calculation tools. Principal Component Analyses (PCA) were then performed using the prcomp function in the R stats package (R Core Team, 2021) for the sites across the six normalised metal axes (Figure 1). Further details on site water chemistry data and sampling regime can be found in Osmond et al., 2024.

### DNA Extraction and Sequencing

DNA was extracted from fin clip samples using a salting out protocol (Jenkins, 2019). DNA quality and quantity was checked by visualisation on a 1% agarose gel and spectrophotometrically using a Nanodrop. Samples were sequenced using DNBseq (BGI, China) with paired-end 150bp reads. Given the broad aims of the questions to be addressed, a combination of sequencing depths was employed. A target coverage of 4x was used for 140 samples, with three samples omitted after failing library preparation. A target coverage of 15x was used for 12 samples to provide a truth-set for high-confidence variant calling, with a further five samples sequenced at 40x to ensure unambiguous alignment of representative metal-impacted individuals with the published brown trout genome. The higher coverage libraries were sequenced on a NovoSeq 6000 (Illumina, USA) with 150 bp paired-end reads. Full details of the sequencing statistics are given in Supplementary Materials 2.

### Quality Control and Variant Calling

Reads were quality checked using FastQC (Andrews, 2010) and were cleaned using TrimGalore (Krueger et al., 2023). Reads were aligned to the *S. trutta* reference genome (fSalTru1.1; Hansen et al., 2021) using BWA mem (Li, 2013) and sorted and indexed using SAMtools v1.3.1 (Danecek et al., 2021). Information on read groups were added, duplicate reads were removed, and samples sequenced across multiple lanes were merged using picard v2.6.0 (“Picard Tools.,” 2016). Variants were called using both GATK4 (Van der

Auwera and O’Connor, 2020) and Freebayes v1.3.1 (Garrison and Marth, 2012) across the 17 15-40x individuals to generate a high confidence set of variant sites. GATK-called SNPs were filtered according to GATK best practices. Both Freebayes-SNPs and GATK-SNPs were filtered using vcftools v0.1.15 (Danecek et al., 2011), retaining only biallelic SNPs with a minimum minor allele frequency (MAF) of ≥0.05, a maximum missingness of 10% per population, a minimum quality score of 30 and a read depth between 5x and 140x. Filtered sites were intersected using bcftools v1.9 (Danecek et al., 2011), resulting in a final set of 5,784,552 high-confidence biallelic SNPs, of which 4,875,231 mapped to chromosomal contigs and were retained for selection scans.

### Calling genotype likelihoods and population-level genomic diversity

Genotype likelihood files were generated using ANGSD (Korneliussen et al., 2014), using the high-confidence variants as a SNP list, for the 137 samples sequenced at 4x and the twelve 15x samples. Likelihoods were calculated with the ANGSD genotype likelihood (GL) utility, using the SAMTools GL model and with the reference genome allele fixed as the major allele, to produce genotype likelihoods for 149 individuals. Custom scripts (see Data Availability) were used to calculate the minor allele frequency (MAF) for each sampled population, filtering sites which had a MAF<0.05 across the low coverage individuals.

Site allele frequency (saf) likelihoods were calculated for each population in ANGSD using the –dosaf command with the GATK genotype likelihood model and the ancestral population set as the reference genome allele. The realSFS module in ANGSD was applied to the saf file of each population to produce one-dimensional site frequency spectrum (SFS) files. Index files were produced between population pairs, two dimensional SFS calculated, and global estimates of *F_ST_* were calculated using the realSFS fst stats function.

### Estimating evolutionary relationships between populations

Treemix v1.13 (Pickrell and Pritchard, 2012) was used to contextualise genetic relationships between populations, rooted on NEV due to having the lowest average pairwise *F_ST_*with other sampled populations and being relatively unimpacted by minewater pollution. Unrooted analyses were run initially alongside analyses but were not informative. To estimate the optimal number of migration edges, 100 runs were conducted for *m = 1-10*, with a SNP block size of 1000, with results compared using the Δm Evanno method implemented in the optM package in R (Fitak, 2021). 500 independent runs were conducted at the optimal level of m. The best likelihood tree was retained, bootstrap support estimated using Phylip Consense v3.697 (Felsenstein, 2004) and output plotted using BITEV2 package (Milanesi et al., 2017). Output figures for optimal migration edge number selection are given in Supplementary Material 1 Figure S3.

Each chromosome was split into a Beagle file format (Browning, 2018) and was subsequently converted into .pos format using custom scripts in Unix. Resulting files were used as input for ngsLD (Fox et al., 2019) to calculate linkage between loci on each contig using default parameters. Linked loci were pruned with maximum distance of 2000 kb and minimum weighting of 0.5. The genotype probability format file was then filtered to retain only unlinked loci. Unlinked loci were used to perform individual-based Principal Component Analysis (PCA) using PCAngsd with default parameters. Resulting eigenvectors and eigenvalues were plotted in R using PCAngsd scripts (Meisner et al., 2021).

To estimate genome-wide nucleotide diversity across populations, the thetaStat tool from ANGSD was used to calculate statistics of Pi and Tajima’s D in windows of 10 kb within each population from the folded SFS. Genome-wide and plots per population were plotted in R using ggplot2 (Wickham, 2016). The significance of the effect of metal-impact on nucleotide diversity and Tajima’s D was tested utilising linear mixed effects models in the lme4 package (Bates et al., 2015), with the individual population fitted as a random effect to account for individual variance between populations. Significance of metal impact was tested using factor reduction and likelihood ratio test.

### Identifying genomic regions of parallel selection

Filtered allele frequency estimates for each individual population were used as an input matrix to AF-vapeR (Whiting et al., 2022). Population vectors were defined at a number of levels to investigate parallelism: 1) across sampling sites comprising all sampled regions of the British Isles; 2) within regions where multiple sites have experienced the same stressor; 3) within rivers experiencing relatively similar metal pollution (Figure 1). Details of the populations used in each vector set are given in Table 3. Vectors were defined to maximise the gradient between the metal-impacted and clean control populations and to balance the number of vectors within each region to reduce signals of phylogeographic bias. Analysis was run individually across each of the 40 chromosomes. Allele frequency change vectors were calculated in windows of 200 SNPs with 1000 null permutations per chromosome. Eigen analysis was performed for the allele frequency change vectors and null permutations and p-values were calculated using the eigen_pvals function. Significant windows were identified using the signif_eigen_windows function, using null permutation cut-off values of >99% and 99.9%. Results were plotted in R using ggplot2 (Wickham, 2016).

**Table 3.** Population vectors applied for the AF-vapeR analyses. Five separate analyses were applied to investigate parallel adaptation across the full geographic range (Britain and Ireland), and within local regions of Wales and NE England, and within predominantly copper-impacted and lead-impacted catchments. Each population pair in the analysis is represented by a metal-impacted and a relatively clean control population.

| Vector | Region | Target population | Control |
| --- | --- | --- | --- |
| **Britain and Ireland** | Anglesey | AGA | AGB |
|  | Ireland | AVB | OWE |
|  | Cornwall | HAY1 | CAM |
|  | Mid Wales | YHE | NEV2 |
|  | North Wales | CWA | CWC |
|  | NE England 1 | TYD | TYA |
|  | NE England 2 | WEA | WEF |
| **Wales** | Anglesey | AGA | AGB |
|  | Mid Wales | YHE | NEV2 |
|  | North Wales 1 | CWA | CWC |
|  | North Wales 2 | CWB | CWC |
| **NE England** | Tyne 1 | TYD | TYA |
|  | Tyne 2 | TYE | TYA |
|  | Wear | WEA | WEF |
| **Copper** | Anglesey | AGA | AGB |
|  | Ireland | AVB | OWE |
|  | Cornwall | HAY1 | CAM |
| **Lead** | Tyne 1 | TYD | TYA |
|  | Wear | WEA | WEF |
|  | Mid Wales | YHE | NEV2 |
|  | North Wales | CWA | CWC |

We also applied PicMin (Booker et al., 2023) to detect parallel adaptation. Pairwise weighted *F_ST_* analysis between each of the population pairs defined previously using AF-vapeR was conducted in non-overlapping windows of 50 kb using the realSFS fst stats2 function. This window size was chosen to reduce the number of statistical comparisons and thus the FDR scaling, whilst still providing sufficient specificity to detect regions under selection. Windows not overlapping with Ensembl annotated genes from the *S. trutta* genome were removed using the GenomicRanges package in R (Lawrence et al., 2013) resulting in 33,500 gene-overlapping windows. Empirical *F_ST_* p-values for each window in each population pair were calculated using the empPvals function from the qvalue package in R (Storey et al., 2023). Tippett nulls were calculated for the number of population pairs for each analysis with 1,000,000 repetitions. PicMin analysis was run to calculate a p-value of significance for each window of the genome with the Tippett nulls used as the null values. FDR adjustment of p-values was performed using the p.adjust function with the FDR method used for correcting multiple comparisons. Populations contributing to a significant window were extracted and plotted using the pheatmap package (https://CRAN.R-project.org/package=pheatmap) in R (Supplementary Materials 1, Figure S2). Genes overlapping significant windows were annotated using the BioMart package in R (Smedley et al., 2009).

PCA-based methods integrating a probabilistic framework have been suggested as a powerful method of detecting selection from low-coverage whole-genome data (Lou et al., 2020). We therefore used the selection utility of PCAngsd (Meisner et al., 2021), which utilises fastPCA whilst accounting for genotype uncertainty. We applied this within three groups: i) all individuals in the data set, and separate runs including only those sampling sites representing ii) northeast England, and iii) Wales. Genotype probabilities were used as input, filtering for MAF < 0.05. Resulting covariance matrices and selection statistics for each run of PCAngsd selection were used as input into R for plotting, using custom scripts adapted from those provided in the PCAngsd public GitHub repository (https://github.com/Rosemeis/pcangsd/tree/master). Test statistics were converted to p-values and quantile-quantile (QQ) plots were generated to ensure that results followed the assumed chi-squared distribution. The position of variants used in each analysis and associated p-values along the examined PC axes were combined into a matrix and exported for downstream analysis. Significant outlier loci were extracted along each PC axis where p<1e^-7^. Variants identified within 10 kb and overlapping with the same gene annotation were merged. Manhattan plots of –log(p) were plotted for each PC axis in R using custom scripts and ggplot2 (Wickham, 2016).

### Identifying unique genomic regions of adaptation within populations

To investigate evidence of population-specific evolutionary responses, we applied population branch statistics (PBS) (Yi et al., 2010). The population groupings used for the PBS analyses can be found in Table 4, with control populations selected on the basis of relatively low metal-impact and relatively low genome-wide average *F_ST_* between populations. The three focal populations were indexed together with realSFS, and PBS statistics were calculated using the *fst stats2* option with non-overlapping windows of 50 kb. Empirical p-values for PBS values of each population were calculated using the empPvals function in the *qvalue* package in R (Storey et al., 2023). These were combined into a matrix of chromosome, sequence window and p-value for each focal population. To identify putative outlier windows, we selected windows with an empPval < 0.005 in at least one of the population groups. Genes overlapping PBS outlier windows were annotated using the GenomicRanges (Lawrence et al., 2013) and biomaRt packages in R (Smedley et al., 2009).

**Table 4.** Population groupings used in the PBS analyses, with the group name for the analysis, the focal metal-impacted population, paired clean-control population and outgroup clean population.

| Group Name | Focal Population | Paired Clean | Outgroup Clean |
| --- | --- | --- | --- |
| Tyne_1 | TYD | TYA | WEF |
| Tyne_2 | TYE | TYA | WEF |
| Wear | WEA | WEF | TYA |
| Anglesey | AGA | AGB | NEV |
| Cornwall | HAY | CAM | NEV |
| Ystwyth | YHE | NEV | CAM |
| Avoca | AVB | AVD | OWE |
| Conwy_1 | CWA | CWC | NEV |
| Conwy_2 | CWB | CWC | NEV |

We utilised Sweepfinder2 (DeGiorgio et al., 2016) to identify regions of the genome that differed from the null expectation from a genome-wide site frequency spectrum (SFS) within each of our putatively metal-adapted populations. Allele frequency tables per population were converted into the required .plink.freq file format using custom scripts in R. A global estimate of the SFS per population was calculated using SweepFinder2 prior to outlier analysis. A recombination map for brown trout mapped to the current reference genome was unavailable and a B-map was therefore not used to control for the confounding signals of background selection. A predefined distance of 5 kb was applied and composite likelihood ratios (CLR) were calculated for each population. Any outlier windows within 100 kb of each other were merged to be considered as selection acting on the same genomic region and non-overlapping windows with paired-control populations were used to remove spurious signal from background selection.

### Evaluating concordance between different methods of selection outlier analysis

To examine concordance between the various methods for detecting regions under selection, the GenomicRanges (Lawrence et al., 2013) package was used to identify overlapping regions identified by the three methods applied to identify parallel adaptation (AF-vapeR, PCAngsd and PicMin). Genomic regions identified as significant above the 99^th^ percentile of null permutations were extracted from each of the AF-vapeR vector set analyses and overlapped with the PicMin FDR<0.5 outliers, filtering these for only those windows where the empPval for all of the focal populations for that vector set were below 0.05. For AF-vapeR vector sets which represented just a subset of populations, either by water chemistry or region, the overlap was subsequently filtered to retain only windows where the empPval of *F_ST_* in the PicMin analysis was less than 0.05 for each population pair used in the vector set. These AF-vapeR windows were then subsequently filtered to retain only windows that had no antiparallel populations along the first eigenvector in the AF-vapeR analysis. Outliers from each method and the intersected genes common to both AF-vapeR and PicMin were also intersected with the candidate outlier loci for the NE England and Wales PCAngsd analyses.

To find concordance in methods targeting individual adaptations in each target population, we used the GenomicRanges package in R (Lawrence et al., 2013) to subset the SweepFinder2 outliers above the 99.5 percentile with the PBS outlier windows with an empPvalue of < 0.01 for each focal population.

### Intersection with gene expression evidence

We intersected selection outliers from our dataset with constitutively overexpressed genes (logFC >1, FDR ≤ 0.05) identified in the gill (n=2,042) and liver (n=311) of metal-tolerant Hayle trout following depuration (Paris et al., 2024), anticipating that genes maintained under both differential expression and positive selection are most likely to represent candidates underlying an adaptive response to metal toxicity.

Our candidate regions were intersected in R (R Core Team, 2021) by combining the unique Ensembl gene identifiers of the expression outliers and the genes overlapping outlier windows from each method we applied to identify selection outliers. Outlier regions for parallel methods were defined as FDR < 0.5 for PicMin, and fully-parallel windows above 99% of null permutations in the AF-vapeR analyses. We also intersected the SweepFinder2 CLR and PBS outlier windows and Hayle PBS windows found to have ap-value < 0.005. For each method, two lists were produced for overlapping genes for the expression outliers from the gills and for the liver. We then applied the reduce function from base R (R Core Team, 2021) with the intersect option, to identify genes present in both the selection outlier scans and constitutively expressed in the transcriptomics study.

### Read Depth Mapping

For visualisation of potential deletions or duplications driving the genomic divergence identified by SNP markers within the chr25 region, read depth of medium-high depth sequenced individuals was investigated. Supplementary alignments were first removed from bam files using SAMtools v1.9 (Danecek et al., 2021) before depth per base along the *S. trutta* reference genome was calculated with SAMtools depth. These were subset to retain the focal area of interest and plotted in R (R Core Team, 2021), utilising ggplot2 (Wickham, 2016) and overlapping genes within the region labelled with gggenes (Wilkins, 2020).

### Gene Ontology Enrichment analysis

Outlier windows identified by the selection tests were examined using Gene Ontology (GO) enrichment to test for significantly overrepresented Biological Process (BP) and Molecular Function (MF) terms represented by the identified candidate genes. Analysis was performed in R using clusterProfiler (Wu et al., 2021). Enrichment was conducted using a FDR correction of p-values, GSSize (gene-size) set between 5 and 500, and a q-value cut-off of 0.2.Lists containing GO categorisation of brown trout terms (AH85410) were downloaded from Ensembl using BioMart v2.44 (Smedley et al., 2009). FDR-adjusted p-value cut-offs were at 0.05 for PicMin outlier gene analyses, and 0.1 for PBS-SweepFinder intersected genes, AF-vapeR, and AF-vapeR and PBS intersected genes.

## Supplementary Material

Supplementary Materials 1:

Figure S1 - Manhatten plots of Sweepfinder and PBS outliers.

Figure S2: Heatmap plotting of shared windows between populations in the PicMin FDR < 0.5 outliers.

Figure S3: Treemix test statistics plotting.

Supplementary Materials 2: Sample and sequencing information.

Distribution of sequencing effort across sites and regions. Number of individual unique samples from which DNA was extracted and submitted for sequencing is given in the final three columns.

Supplementary Materials 3: Table of all significant candidate loci for adaptation to metal-enriched environments. Genes overlapping candidate regions are listed, alongside focal population(s), analysis method and level of significance.

## Author Contributions

DRO carried out the lab work, data analysis and wrote the manuscript.

JRP assisted with the study design, data analysis and improved the manuscript.

JFO assisted with data analysis, visualisation and improved the manuscript.

MWB conceived the ideas and experimental design of the study.

JRS conceived the ideas and improved the manuscript.

## Acknowledgements

This research was supported by a NERC GW4 FRESH Doctoral Training Programme studentship. Additional funding was provided by the Game and Wildlife Conservation Trust and an EU Interreg France-England Channel project: Salmonid Management Around the Channel (SAMARCH). Additional funding was provided by the University of Exeter Faculty of Health and Life Sciences Global Lab Visiting Fund to enable collaborative working on the analyses presented here. We are grateful to Dylan Roberts, Dr Charlie Ellis and Greg Wannell for their assistance in collecting samples, and to Emiliano Trucchi and Guillem Izquierdo Aránega for input into the genomic analyses and to James Whiting for input into the running and interpretation of parallel selection methods. We thank Dr Andrew King for comments on earlier drafts of this manuscript. Thanks to Environment Protection Agency Ireland, Natural Resources Wales and The Environment for publicly available water chemistry datasets. Analysis was performed on the High-Performance Computing Cluster (HPCC) (“ISCA”) at the University of Exeter, UK.

## Data Availability

Raw reads for samples contributing to this study will be made available upon publication through the European Nucleotide Archive (ENA). Scripts used for data analysis are available from https://github.com/DanielOsmond/Parallel_Trout/.

