## Supplementary Materials 1 for "Forging together: Parallel adaptation to minewater pollution in brown trout (*Salmo trutta* L.)"

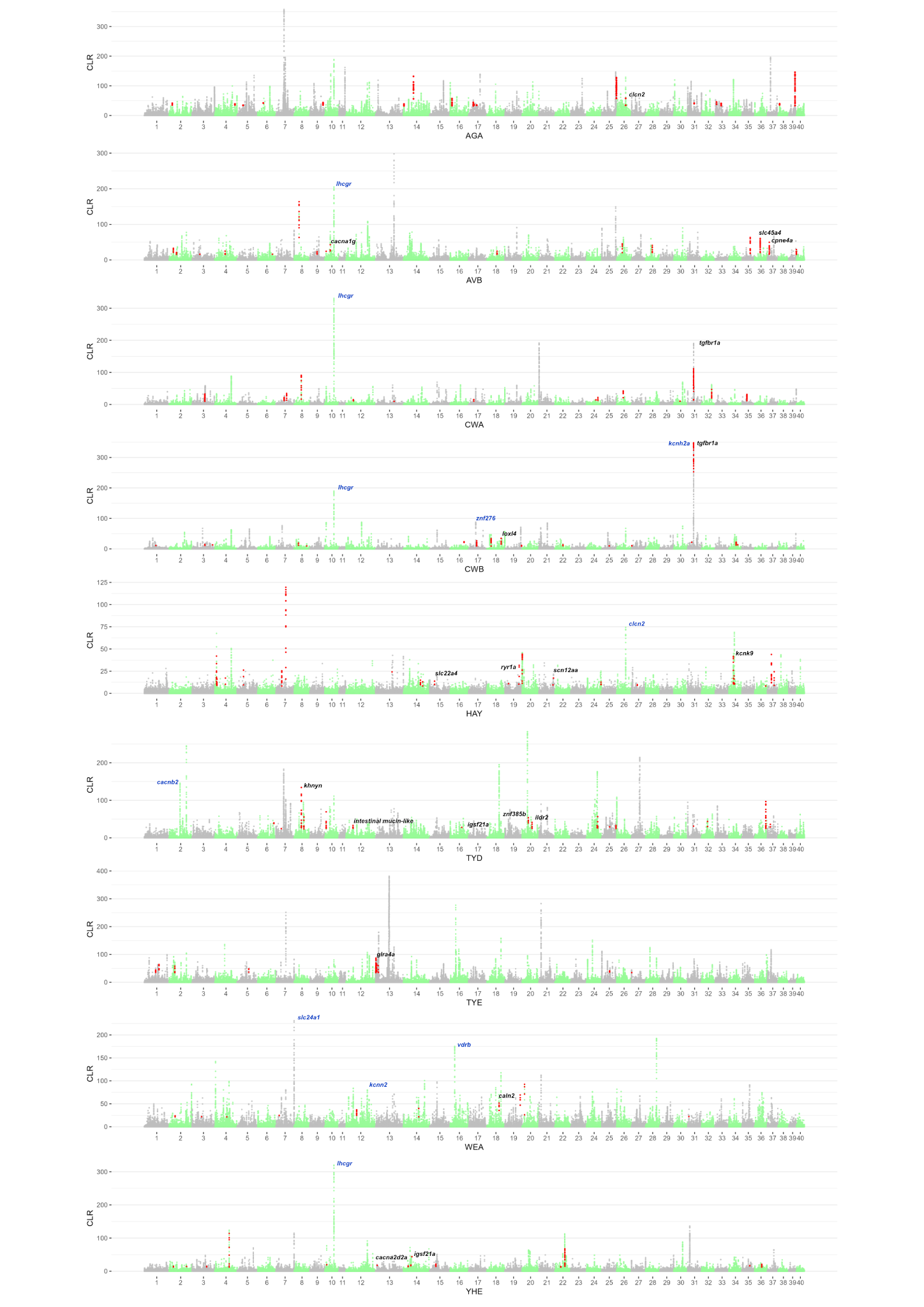

**Figure S1.** Manhattan plots of SweepFinder2 output, based on windows of 5000 bps for each of the focal metal-impacted populations, with composite likelihood ratios (CLR) plotted on the y-axis. Overlaps between the CLR 0.995 percentile outliers and PBS outliers below a p-value of 0.01 are given by red points on the plot. Candidate genes within these overlapping regions are labelled in black, with genes with credible adaptive function but identified by only a single method labelled in blue.

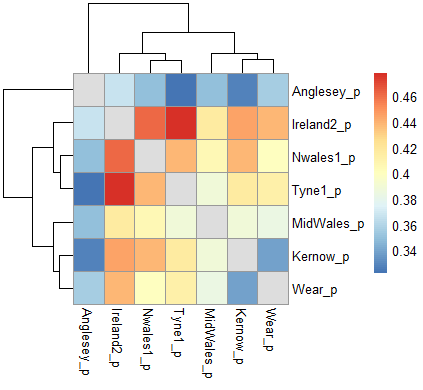

**Figure S2.** Heatmap of shared windows between populations contributing towards 170 PicMin outlier windows at FDR < 0.5. The relatively balanced proportion of shared windows across population vectors suggests that over-representation of geographically proximate populations is not biasing analysis.

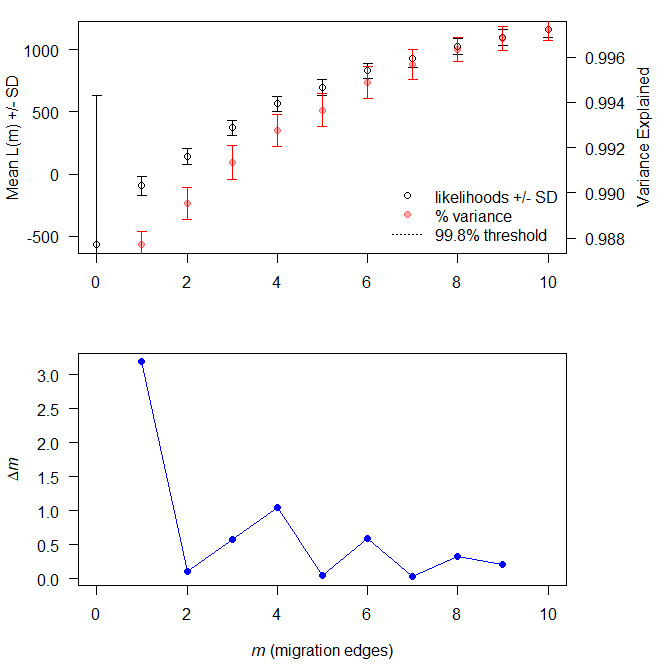

**Figure S3.** Test statistics plots for Treemix runs to determine optimum level of migration edges. Top: The mean and standard deviation (SD) across 10 iterations for the composite likelihood *L*(*m*) (left axis, black circles) and proportion of variance explained (right axis, red “x”s). Bottom: The second-order rate of change (Δm) across values of m in sampled trout populations.
